# Knocking-out the human face genes *TBX15* and *PAX1* in mice alters facial and other physical morphology

**DOI:** 10.1101/2021.05.26.445773

**Authors:** Yu Qian, Ziyi Xiong, Yi Li, Haibo Zhou, Manfred Kayser, Lei Liu, Fan Liu

**Affiliations:** CAS Key Laboratory of Genomic and Precision Medicine, Beijing Institute of Genomics, Chinese Academy of Sciences, Beijing, China; China National Center for Bioinformation, Beijing, China; University of Chinese Academy of Sciences, Beijing, China; Department of Genetic Identification, Erasmus MC University Medical Center Rotterdam, Rotterdam, the Netherlands; Department of Epidemiology, Erasmus MC University Medical Center Rotterdam, Rotterdam, the Netherlands; Institute of Neuroscience, State Key Laboratory of Neuroscience, Key Laboratory of Primate Neurobiology, CAS Center for Excellence in Brain Science and Intelligence Technology, Shanghai Institutes for Biological Sciences, Chinese Academy of Sciences, Shanghai, China; Department of Plastic and Burn Surgery, The Second Hospital, Cheeloo College of Medicine, Shandong University, Jinan, China

## Abstract

DNA variants in or closed to the human *TBX15* and *PAX1* genes have been repeatedly associated with facial morphology in independent genome-wide association studies, while their functional roles in determining facial morphology remains to be understood. We generated *Tbx15* knockout (*Tbx15^-/-^*) and *Pax1* knockout (*Pax1^-/-^*) mice by applying the one-step CRISPR/Cas9 method. A total of 75 adult mice were used for subsequent phenotype analysis, including 38 *Tbx15* mice (10 homozygous *Tbx15^-/-^*, 18 heterozygous *Tbx15*^+/-^, 10 wild-type WT) and 37 *Pax1* mice (12 homozygous *Pax1^-/-^*, 15 heterozygous *Pax1*^+/-^, 10 WT mice). Facial and other physical morphological phenotypes were obtained from three-dimensional (3D) images acquired with the HandySCAN BLACK scanner. Compared to WT mice, the *Tbx15^-/-^* mutant mice had significantly shorter faces (*P*=1.08E-8, R2=0.61) and their ears were in a significantly lower position (*P*=3.54E-8, R2=0.62) manifesting an “ear dropping” characteristic. Besides these face alternations, *Tbx15^-/-^* mutant mice displayed significantly lower weight as well as shorter body and limb length. *Pax1^-/-^* mutant mice showed significantly longer noses (*P*=1.14E-5, R2=0.46) relative to WT mice, but otherwise displayed less obvious morphological alterations than *Tbx15^-/-^* mutant mice did. Because the *Tbx15* and *Pax1* effects on facial morphology we revealed here in mice are largely consistent with previously reported *TBX15* and *PAX1* face associations in humans, we suggest that the functional role these two genes play on determining the face of mice is similar to the functional impact their human homologues have on the face of humans.

**Author Summary:** Several independent genome-wide association studies (GWASs) on human facial morphology highlighted DNA variants in or closed to *TBX15* and *PAX1* with genome-wide significant association with human facial phenotypes. However, direct evidence on the functional involvement if these genes in the development and determination of facial morphology has not been established as of yet. In the current study, our *in vivo* gene editing experiments in mice for two well-replicated human face *TBX15* and *PAX1* genes provide novel evidence on the functional involvement of these two genes in facial and other physical morphology in mice, at least. *Tbx15^-/-^* mice showed a shortened facial length and manifesting an ear dropping characteristic, *Pax1^-/-^* mice showed an increased nose length. Our geometric morphometrics analysis further indicate that there are significant facial morphology differences between groups (*Tbx15^-/-^*and *Tbx15^+/-^, Tbx15^-/-^* and *Tbx15^+/+^, Tbx15^+/-^* and *Tbx15^+/+^, Pax1^-/-^* and *Pax1^+/+^*). We provide the first direct functional evidence that two well-known and replicated human face genes, *Tbx15* and *Pax1*, impact facial and other body morphology in mice. The general agreement between our findings in knock-out mice with those from previous GWASs suggests that the functional evidence we established here in mice may also be relevant in humans.

## Introduction

Human facial morphology represents a set of highly variable, multidimensional, highly correlated, symmetrical, and strongly heritable complex phenotypes. Unveiling the genetic basis of human facial variation is of fundamental and applied value in developmental biology, evolutionary biology, human genetics, medical genetics, and forensic genetics.

Several independent genome-wide association studies (GWASs) on human facial morphology highlighted DNA variants in or closed to *TBX15* and *PAX1* with genome-wide significant association with human facial phenotypes [1-6]. In particular, Xiong et al. found DNA variants near *TBX15* and *PAX1* significantly associated with human face length and nose width, respectively [1]. Adhikari et al. reported SNPs in *TBX15* to be significantly associated with two human ear traits, folding of antihelix and antitragus size [2]. Claes et al. and White et al. showed *TBX15* and *PAX1* intron variants to be significantly associated with human facial shape [3,6]. Adhikari et al. and Shaffer et al. reported significant association of *PAX1* intergenic variants with human nose wing breadth [4,5].

*TBX15* is expressed in limb mesenchymal cells and has a significant function in limb development [7-9]. Genetic variants in *TBX15* result in cousin syndrome, characterized by craniofacial deformity, scapular hypoplasia, pelvic dysplasia, short stature [10-12]. *PAX1* mutations cause autosomal recessively inherited otofaciocervical syndrome, characterized by facial dysmorphism and external ear anomalies [13]. *PAX1* acts as an important regulator of chondrocyte maturation [14]. Deletion of the short arm region of chromosome 20 including the *PAX1* gene locus cause craniofacial deformities and abnormal vertebral bodies [15].

Taking all available genetic knowledge together allows concluding that *TBX15* and *PAX1* play a role in facial morphology in humans. However, direct evidence on the functional involvement if these genes in the development and determination of facial morphology has not been established as of yet. Although previous *in vivo* experiments revealed that *Tbx15^-/-^*mutant mice exhibited lower weight, glucose intolerance, and obesity on high fat diets [16-18], these studies did not explore the effect of *Tbx15* on facial morphology in mice. Moreover, no functional work has been reported for *Pax1* in mice, or other functional evidence for *PAX1* in humans.

In this study, to enhance our functional understanding on the involvement of *Tbx15* and *Pax1* in facial morphology in mice, we respectively knocked out these genes by applying the one-step CRISPR/Cas9 method [19]. On mutant and wild-type (WT) mice, we acquired 3D images through the HandySCAN BLACK scanner. Based on the collected digital imagery, we quantitatively assessed facial morphology as pairwise Euclidean distances between a set of anatomically meaningful facial landmarks, and also quantified other morphological phenotypes such as weight, body length, fore and hind limb length. Finally, the morphological differences between mutant mice and WT mice were compared using appropriate statistical tests.

## Results

### Knock-out mice

We generated *Tbx15^-/-^* and *Pax1^-/-^* F0 knock-out mice, respectively, using the one-step CRISPR/cas9 method as described previously (Fig 1A-1B) [19].F1 heterozygotes mice were mated with a female to male ratio of 2:1. After eight months of breeding experiments, we obtained a total of 38 F2 9-weeks adult *Tbx15* mice i.e., 10 homozygous *Tbx15^-/-^*, 18 heterozygous *Tbx15*^+/-^, 10 wild-type *WT*, and 37 F2 9-weeks adult *Pax1* mice i.e., 12 homozygous *Pax1^-/-^*, 15 heterozygous *Pax1*^+/-^, 10 *WT* mice (S3 Table). Genotypes of all mice were confirmed by PCR amplification and subsequent Sanger sequencing (Fig 1C-1D). The birth rate of *Tbx15^-/-^* mice was significantly lower than *WT* mice during a 6-month period (mean birth rate 0.26 vs. 0.74, *P*= 0.04, Fig 2A- 2B), suggesting an important role of *Tbx15* in embryonic development. The birth rate of *Pax1^-/-^* mice was also lower than that of *WT* mice, but not statistically significantly so (Fig 2C-2D).

**Fig 1.**
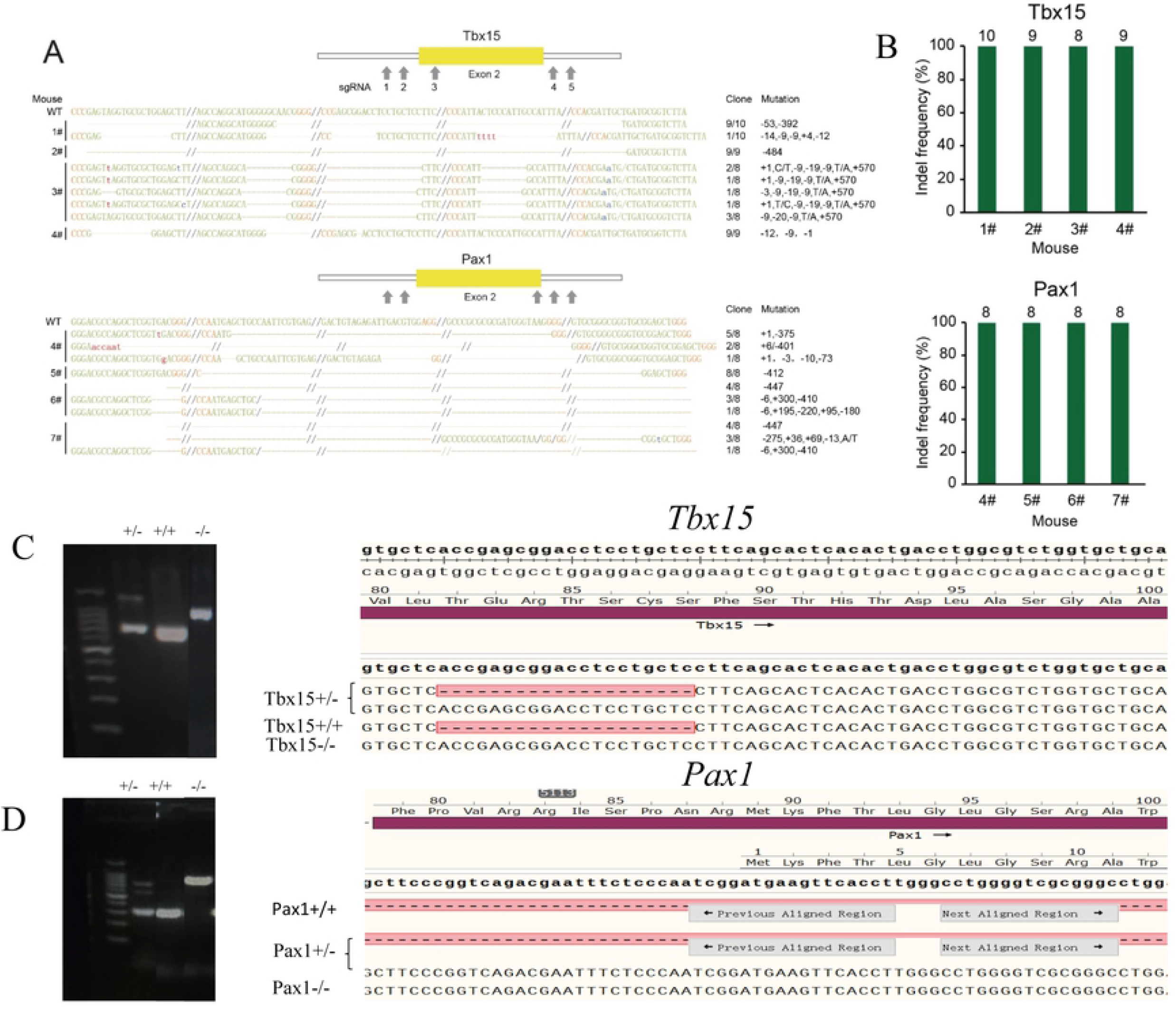
Complete deletion of Tbx15 and Pax1 in mice by CRISPR/Cas9. (A) Clones from tail were sequenced and analyzed. Schematic of sgRNA-targeting sites in Tbx15 and Pax1 gene. The sgRNA target sequences and PAM sequences are labeled in green and red, respectively. Hyphens represent deleted nucleotides and omitted regions are indicated by dash lines. (B)The indel frequencies show that Tbx15 and Pax1 were mutated completely in all mice. Number of clones denoted above the column.(C) Genotype identification for Tbx15 mice by PCR and Sanger sequencing.(D) Genotype identification for Pax1 mice by PCR and Sanger sequencing.

**Fig 2:**
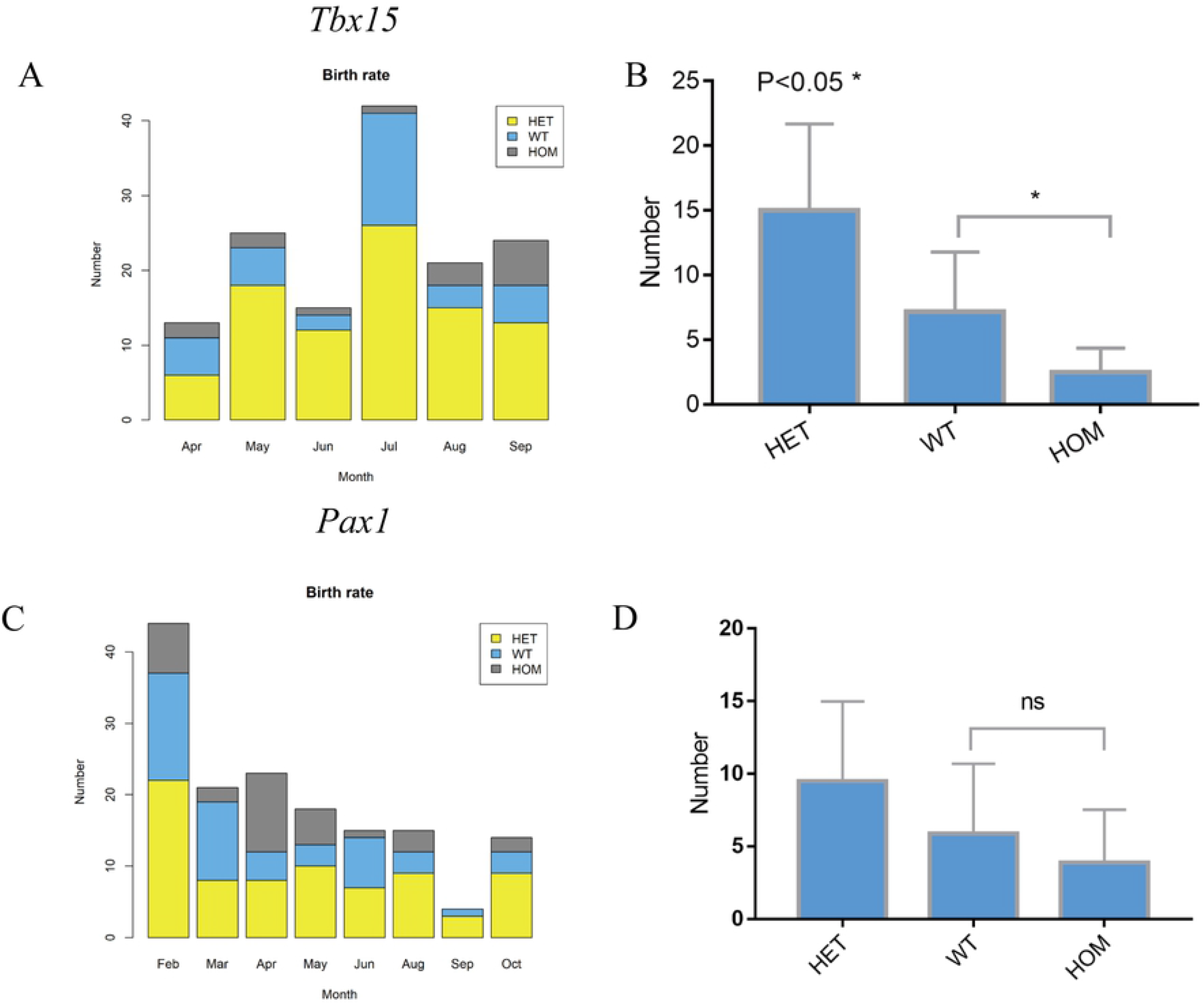
Number of new-born mice with different genotypes. (A) Number of mice with different genotypes in *Tbx15* F2 generation in each month. (B) The t test of the total number of births of *Tbx15* wild-type and homozygous mice. (C) Number of mice with different genotypes in *Pax1* F2 generation in each month. (D) The t test of the total number of births of *Pax1* wild-type and homozygous mice.

### Facial phenotypes

For all 75 F2 mice, digital 3D surface data were collected by use of the HandySCAN BLACK scanner. For quality control purposes, we compared the accuracy of the scanner with a millimeter ruler using a cylindrical object. The height and the diameter measured by the ruler (135.71 and 84.55 mm) were highly consistent with those measured by the scanner (135.70 and 84.58 mm), with the maximal difference smaller than 0.03 mm (S3 Fig). A total of 17 facial landmarks were selected for shape analysis. Among the 17 selected landmarks, 9 overlapped with the 13 facial landmarks in humans as described in the previous face GWAS by Xiong et al (Fig 3A-3B, Fig 4C) [1]. The remaining eight landmarks were all on the mouse ears, and there were no corresponding landmarks in previous GWASs of human face and ear morphology. A total of 136 Euclidean distances connecting all possible pairs obtainable from the 17 facial landmarks were estimated and considered as facial phenotypes in subsequent analyses (Fig 3A-3B). In all WT mice, the facial phenotypes were largely normally distributed (S4 Table), and no significant differences between males and females were observed (S5 Table). As expected, symmetric facial phenotypes showed higher correlations than non-symmetric ones (S6 Table), further supporting the reliability of the obtained phenotype dataset.

**Fig 3.**
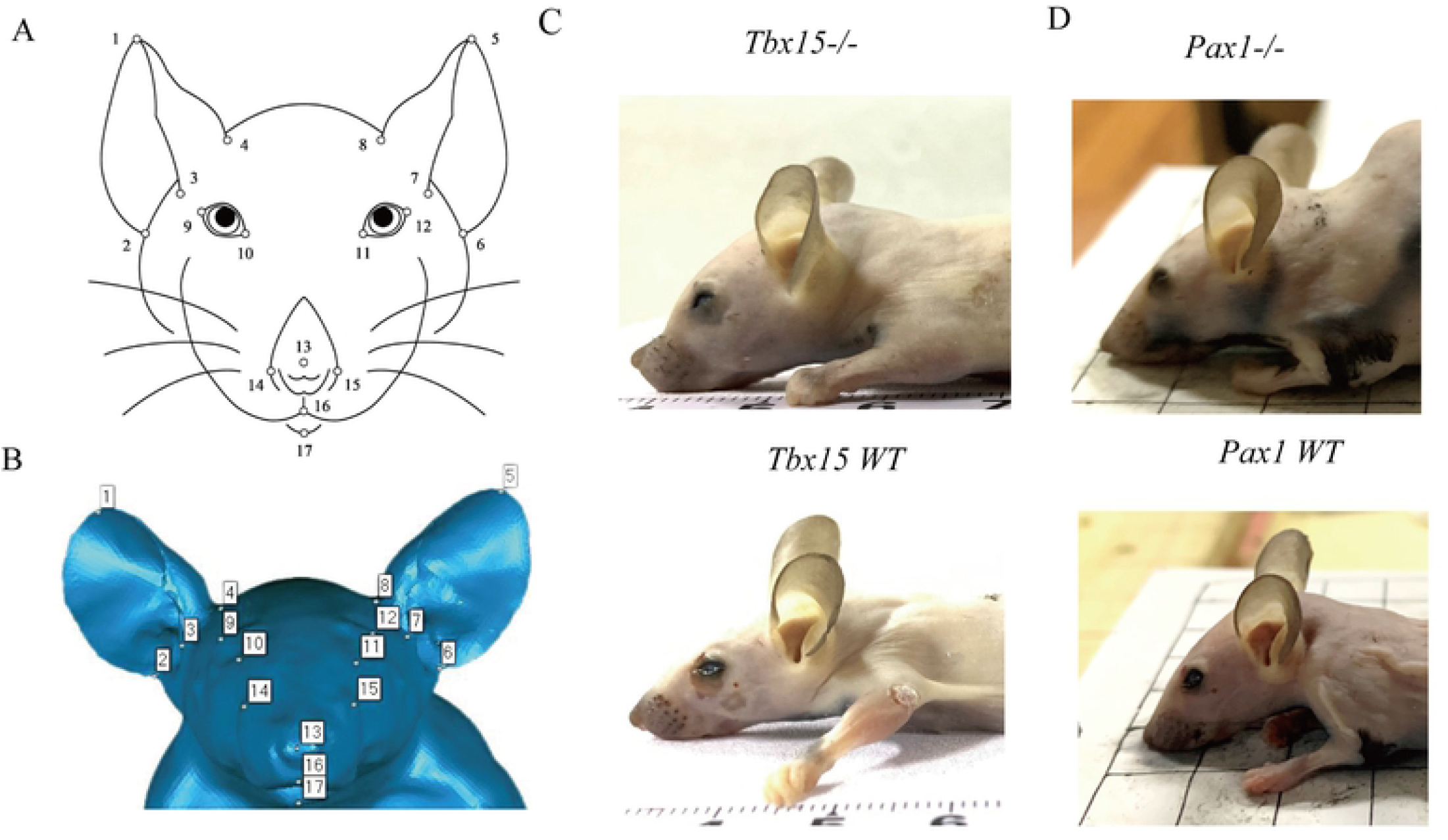
Images of mice. (A) Two-dimensional image of mice. (B) three-dimensional punctuation image of mice. (C) Two-dimensional image of *Tbx15* mice. (D) Two-dimensional image of *Pax1* mice.

**Fig 4.**
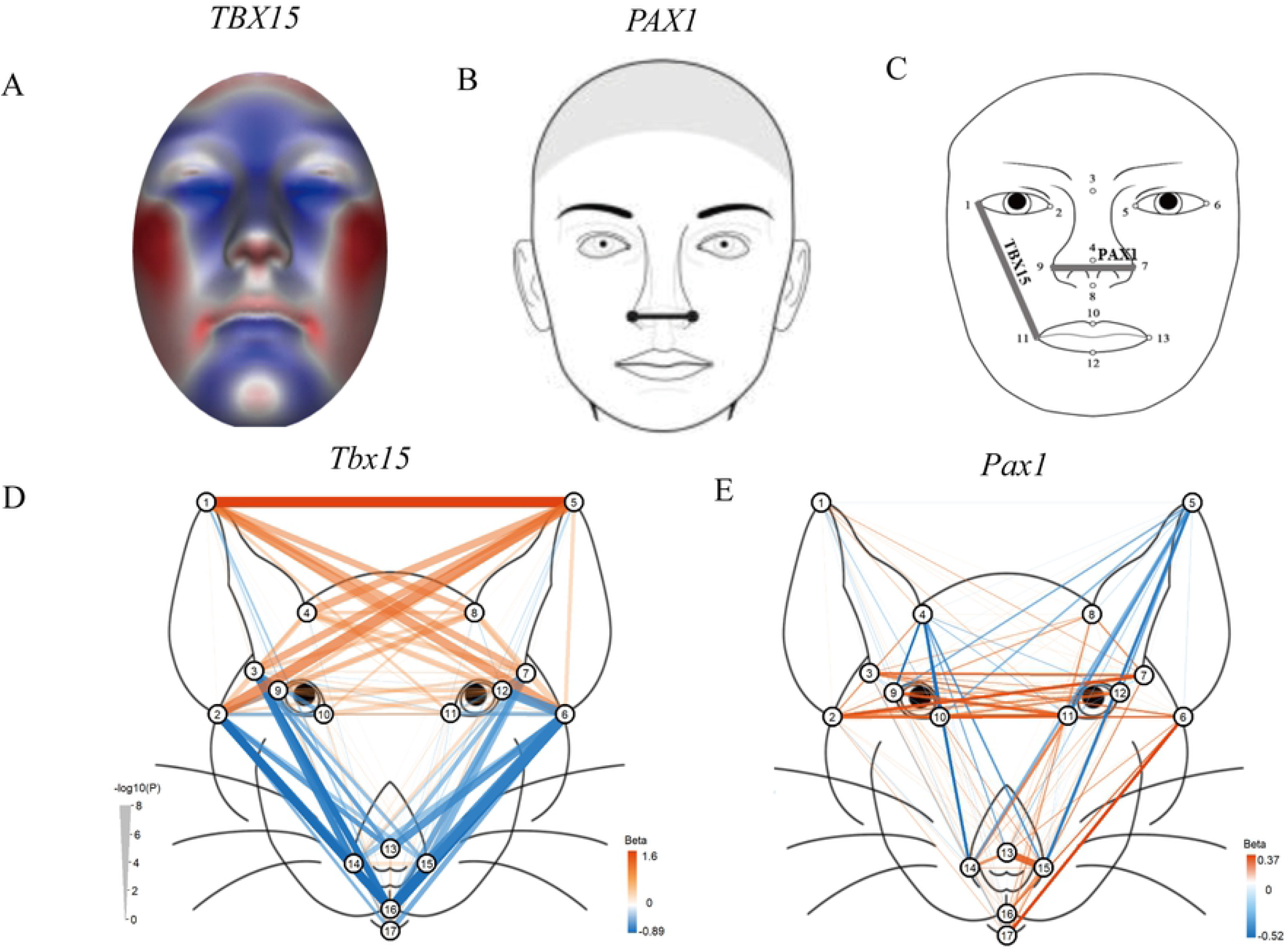
Comparison of mice facial morphology with human. (A) The effect of *TBX15* on human facial morphology[6]. (B) The effect of *PAX1* on human facial morphology[21]. (C) The effect of *TBX15 and PAX1* on human facial morphology[1]. (D) The effect of *Tbx15* on mice facial morphology. (E) The effect of *Pax1* on mice facial morphology

### Differences in facial phenotypes between mutant and wild-type mice

In the 38 Tbx15 mice i.e., 10 homozygous Tbx15^-/-^, 18 heterozygous Tbx15^+/-^, 10 WT *Tbx15^+/+^*, the Tbx15 genotype was significantly associated with facial variation for a total of 34 phenotypes after Bonferroni correction of multiple tests (adjusted P < 0.05). Compared with their WT counterparts, the Tbx15^-/-^mutant mice had significantly shorter faces, which affected 22 phenotypes as characterized by shorter distances between ear root and nose, mouth landmarks, and their ears had a significantly lower position, manifesting an “ear dropping” characteristic. (12 phenotypes, including L1-L8, Fig 3C and 4D, S7 Table). The most significant face shortening effect in the Tbx15 mutant mice was observed for L6-L15, for which the Tbx15 genotype accounted for 61% of the phenotype variance (0.64 mm per mutant gene). The most significant ear dropping effect in the Tbx15 mutant mice was observed for L2-L5 (0.95 mm per mutant gene), with the Tbx15 genotype explaining 62% of the phenotype variance (Figure 4D, S7 Table). Principal components (PCs) were derived from 136 facial Euclidean distance. An unsupervised clustering analysis of PC1 and PC2 could completely separate Tbx15^- /-^ and WT mice (Fig 5A) and a t-test of PC1 demonstrated significant differences between the heterozygote group and the *WT* group (P<.001) as well as between the heterozygote group and the Tbx15^-/-^ group (P<.001, Fig 5B). These results overall demonstrated significant facial differences among Tbx15^-/-^, Tbx15^+/-^ and *WT*.

**Fig 5.**
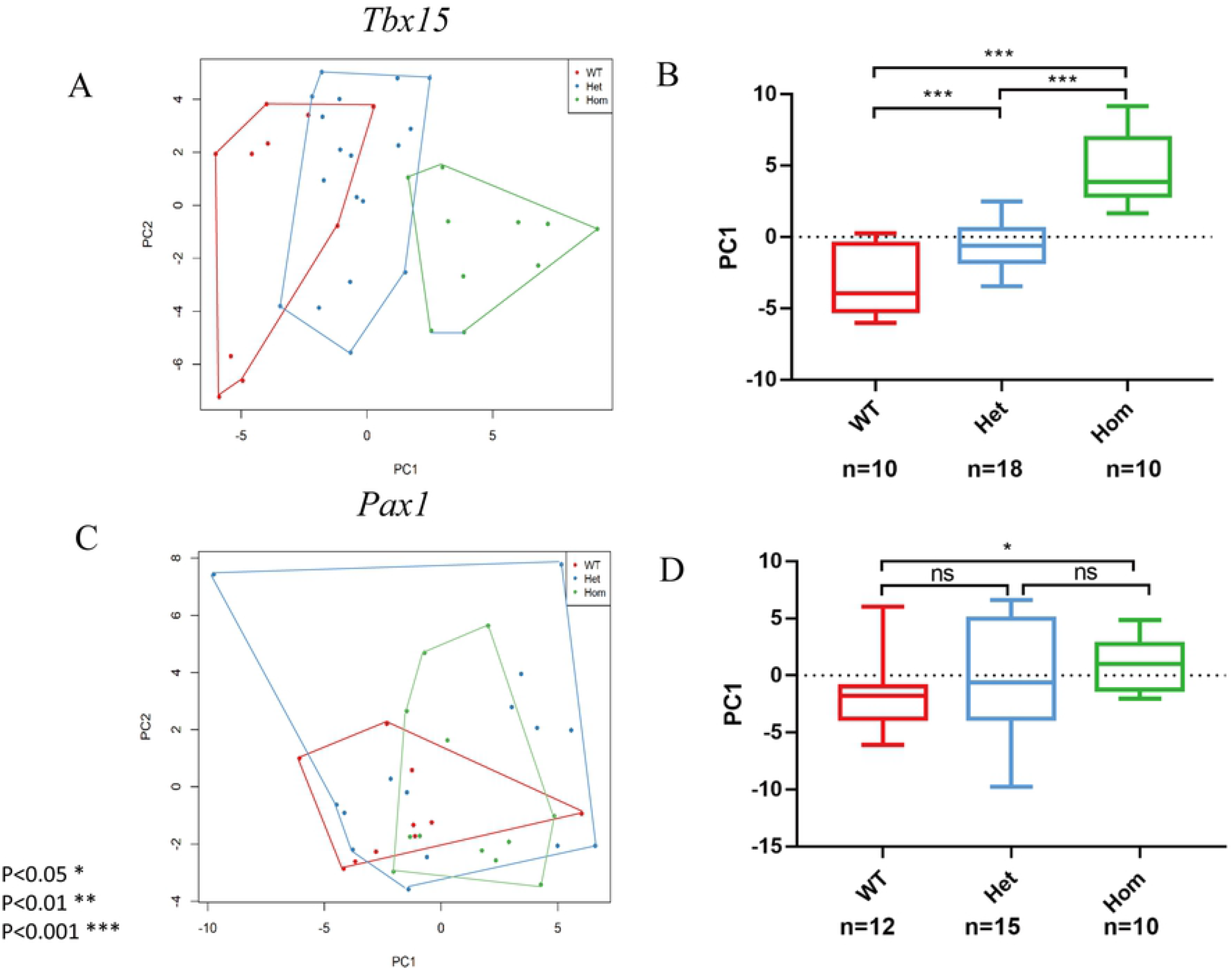
Genotypes classification based on principal components (PCs) of 136 facial distances. (A) Genotypes classification in *Tbx15* mice. (B) PC1 differences between *Tbx15* groups. (C) Genotypes classification in *Pax1* mice. (D) PC1 differences between *Pax1* groups.

In the 37 *Pax1* mice, i.e., 12 homozygous *Pax1^-/-^*, 15 heterozygous *Pax1*^+/-^, 10 WT mice *Pax1^+/+^*, the *Pax1* genotype was significantly associated with facial variation for only one phenotype after Bonferroni correction (adjusted *P* < 0.05). Compared with the WT mice, the nose length of the *Pax1^-/-^* mutant mice were significantly increased (L13-L15, Fig 3D and 4E, S8 Table). The *Pax1* genotype explained 46% of the phenotype variance in the *Pax1* mutant mice (0.28 mm per mutant gene).The clustering analysis of the major PCs could not separate *WT, Pax1^+/-^*, and *Pax1^-/-^* groups. However, the t-test of PC1 showed significant difference between the *WT* and *Pax1^-/-^* groups (Fig 5C-5D).

For both genes, the trends of the facial differences observed between mutant and WT mice were largely consistent with the genetic association effects observed in previous human GWASs (Fig 4) [1-6,20,21]. In particular, *TBX15* variants previously showed significant association effect on multiple facial phenotypes in humans (Fig 4A) [6] and *Tbx15* mutant mice had shorter faces as demonstrated here. The ear dropping characteristic, however, which we observed in *Tbx15* mutant mice had no correspondence with findings in previous human GWASs. Interestingly, Xiong et al. found that SNPs at *TBX15* had asymmetrical effects on human facial morphology (Fig 4C) [1]. In our *in vivo* mice experiments, *Tbx15^-/-^* mice showed highly symmetrical effects on decreasing ear-mouth distances (1.00E-8<*P*<1.66E-4, Fig 3C and 4D, S7 Table). The failure of observing an asymmetric effect of *Tbx15* in our experiments might be explained by a pronounced effect of gene silencing compared with the potential regulatory effects of the SNPs. For *PAX1*, previous GWASs on facial shape in humans mainly found effects on nose width (Fig 4B-4C) [1,4,5]. In our experimental mice study, we found that *Pax1* mutant mice had a significantly increased nose length (L13-L15, Fig 3D, Fig 4E, S8 Table). No significant changes in nose width were observed in *Pax1^-/-^*mice, which may be due to insufficient sample size and facial structure differences between mice and humans.

### Differences in other physical phenotypes between mutant and wild-type mice

Next to the effects of *Tbx15* and *Pax1* gene editing on facial phenotypes in mice, we additionally investigated the effects on other physical morphological phenotypes such as body weight, body length, and length of fore and hind limbs. Compared with WT mice, *Tbx15^-/-^* mice had significantly lower body weight (p<0.01), shorter body length (p<0.001), and shorter limb length (*P* <0.01), (Fig 6). For *Pax1*, the body weight, body length, limb length of the mutant mice were also reduced compared to WT mice, but not statistically significantly so (S1 Figure).

**Fig 6:**
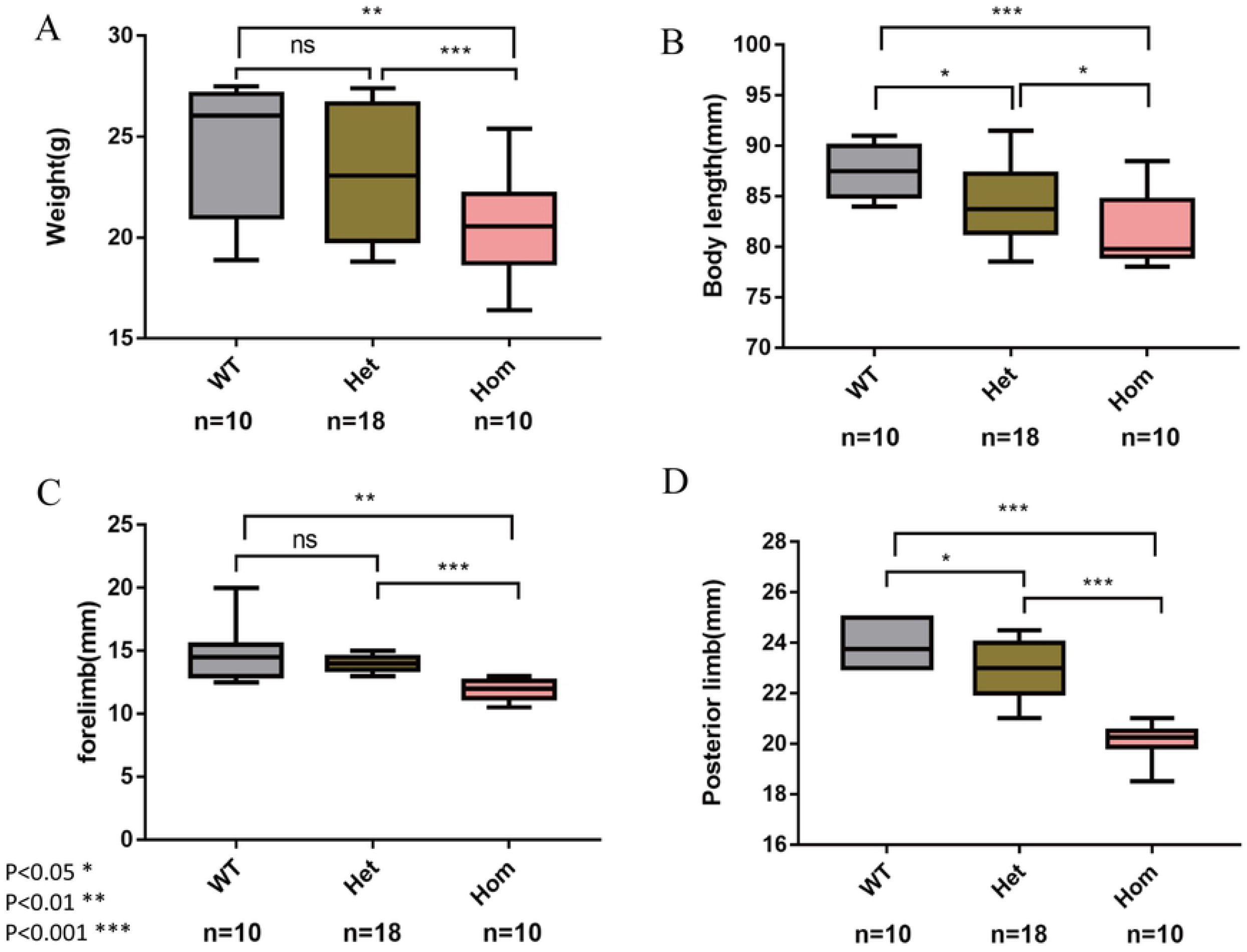
Effects of *Tbx15* on mice weight, body length, fore and hind limb length.

### Protein sequence conservation between mice and humans

The Tbx15 protein sequences are composed of 602 amino acid sequences in both human and mice. An amino acid sequence alignment analysis revealed that the amino acid sequences in both species had very high identity (98.67%, S2A Figure) and the same very long conserved T-box domain was found in humans and mice. The high level of evolutionary conservation between mice and humans, together with the significant lower birth rate of *Tbx15^-/-^* in mice, confirms an important role of *Tbx15* in the early development of embryogenesis, both in humans and in mice.

The Pax1 protein is composed of 534 amino acids sequence in mice and 446 amino acids in humans with a relatively high level of identity (73.41%, S2B Figure), but considerably lower than seen for Tbx15. Furthermore, the two species shared a common set of domains in Pax1, including a ‘Paired box’ domain, a homeodomain-like domain, and a winged helix-turn helix conserved domain. The relatively high sequence conservation may suggest a vital role of *PAX1* in the early development of embryogenesis in both species.

### Expression of *TBX15* and *PAX1* in human embryonic cranial neural crest cells

Embryonic cranial neural crest cells (CNCCs) arise during weeks 3–6 of human gestation from the dorsal part of the neural tube ectoderm and migrate into the branchial arches. The later form the embryonic face, consequently establishing the central plan of facial morphology and determining species-specific and individual facial variation [22,23]. Because the is no expression data in CNCC cells of mice, we took the advantage of the available CNCC RNA-seq data [24] and other public RNA-seq datasets from GTEx [25] and ENCODE [26] in humans and investigated preferential expression patterns in CNCCs. Indeed, both *TBX15* and *PAX1* showed significant preferential expressions in CNCCs compared with that in other cell types (S9 Table). These results indicated important functional roles of these two genes in facial morphogenesis during the early embryogenesis in humans, and likely also in mice.

## Discussion

Our *in vivo* gene editing experiments in mice for two well-replicated human face *TBX15* and *PAX1* genes provide novel evidence on the functional involvement of these two genes in facial and other physical morphology in mice, at least. *Tbx15^-/-^* mice showed a shortened facial length and manifesting an ear dropping characteristic. In addition, they showed reduced weight, shortened body and limb length which are consistent with the findings from a previous mice study [17]. The *TBX15* gene plays a major role in the development of the mesoderm of all vertebrates [27]. The complete inactivation of the mouse *Tbx15* in mice and *TBX15* mutations in human lead to severe bone deformities [8,11,28]. Notably, previous literature also showed that *TBX15* affects the development of head, limbs, vertebrae and ribs by controlling the number of mesenchymal precursor cells and cartilage cells [8], while the *Tbx15* effect on the face was not studied before. Our new findings together with previous ones support our conclusion, that *Tbx15* plays an important role in the development of facial and limb morphology in mice. Because the facial shape effects we observed in the *Tbx15^-/-^* mutant mice are largely in line with the *TBX15* associations effects in humans previously reported in different GWASs, we suggest that the functional impact of *Tbx15* on facial shape we revealed here in mice also exists in humans, which needs to be confirmed by future work.

Moreover, *Pax1^-/-^* mice showed an increased nose length in our study. Earlier studies revealed that *Pax1* is a key transcription factor affecting cartilage development and regulates the expression of cartilage-related genes in the early stages of development [14]. Homozygotes exhibit variably severe morphological alterations of vertebral column, sternum, scapula, skull, and thymus, with reduced adult survival and fertility and some heterozygotes show milder skeletal abnormalities [29-31], while the functional effect of *Pax1* on facial morphology was not studied before. Since the nasal bone contains cartilage component, our findings of *Pax1* determining nose morphology in mice may be explained by its effect on the development of nose cartilage, which needs to be further explored. In our study, we did observe reduced body weight, body length, and limb length in the *Pax1^-/-^* mice compared to WT mice, but the effect was not statistically significant, may be because of small effect and thus insufficient sample size. Although the facial shape effects we observed in the *Pax1^-/-^* mutant mice are not fully consistent with *PAX1* associations effects in humans previously reported in different GWASs, as in humans *PAX1* associations were seen with nose width, while our *in vivo* data show that *Pax1* affects nose length in mice, in both species this gene is involved in nose-related phenotypes. Therefore, we suggest that the functional effect of *Pax1* on facial shape we revealed here in mice is similar to the effect of *PAX1* in humans, which needs to be confirmed by future work.

In addition, these two genes may interact with development-related genes to affect the morphological development*.Tbx15* was co-expressed with *Shox2* genes in multiple tissues in humans and mice [32]. In human, the deletion of the SHOX region on human sex chromosomes caused short stature and *Shox2* plays an important role in the face and body structure formation [33]. Homozygous mice (*Shox2^+/+^*) had been embryonic lethal (63%). The craniofacial development of homozygous in embryonic was slow, and they showed less body weight and shorter body length [29-31]. *Tbx15* and *Shox2* gene may jointly affect the development of the facial and other morphology in humans and mice. There are protein interactions between Pax1 and Meox-1 in humans and mice [34].In human, Meox-1 protein was a mesodermal transcription factor that plays a key role in somitogenesis and was specifically required for sclerotome development. It is required for maintenance of the sclerotome polarity and formation of the cranio-cervical joints [35,36]. In mice, *Meox-1* plays an important role in somatic cell development and limb muscle differentiation. Homozygotes exhibited hemi-vertebrae, rib, vertebral, and cranial-vertebral fusions [29-31].In all, *Tbx15* and *Pax1* may work together with other development-related genes, thus affecting the morphological development in humans and mice, which need to be explored in the future.

In this study, we investigated the function of these two particular human face genes in mice via genes knock-out experiments, because several independent GWASs repeatedly highlighted that intronic and intergenic DNA variants in these two genes demonstrate significant associations with facial shape phenotypes in humans[1-6]. However, functional data to explain these statistical associations was missing thus far and is provided for the first time via direct in-vivo experimental evidence in our present study. In this study, we selected the 9-week-old F2 generation mice because by 9-weeks they become sexually mature before engaging complex behaviors such as fertility, lactation, and reproduction, and their morphological characteristics are well developed. Furthermore, our littermate-born design prevented potential noises caused by unobserved environmental factors. In addition, the inclusion of the heterozygote mice in the analysis further expanded the sample size and was helpful in observing potential trend effects. In addition, we adopted reliable 3D measurement methods. Kaustubh et al analyzed the effect of *EDAR* on ear morphology by acquiring two-dimensional coordinates [2]. Because mice facial phenotypes are consists of facial spatial distances between all combinatorial pairs of the 17 facial landmarks, the three-dimensional method can accurately quantify the appearance of mice and obtains mice 3D images with an accuracy of 0.03mm,which has a wide range of applications in animal morphology study.

This study focused on facial morphology phenotypes in mice, which well-corresponded to the human facial phenotypes investigated in previous GWASs. Future studies may further investigate whether the observed genetic effects can be attributed to bone, fat or muscles.

In conclusion, we provide the first direct functional evidence that two well-known and replicated human face genes, *Tbx15* and *Pax1*, impact facial and other body morphology in mice. *TBX15* seems to affect the face globally, while *PAX1* is mainly affecting the nose, with such a pattern being similar in both mice and humans. The general agreement between our findings in knock-out mice with those from previous GWASs suggests that the functional evidence we established here in mice may also be relevant in humans. In future research studies, functional effects of other human face genes highlighted in previous GWASs should be investigated to take our increasing knowledge on human face genetics from the statistical associational level to the next level of functional proof.

## Materials and methods

### Samples

The mouse strain was C57BL/6J. As the *in vitro* transcription template, Cas9 and sgRNAs were amplified and purified with the gene knockout strategy. Fertilized eggs obtained from super-ovulated females (C57BL/6J, 4 weeks) mated with males (C57BL/6J, 7-8 weeks) were microinjected with mixtures of Cas9 mRNA and sgRNA (Fig 1, S1 Table). The injected fertilized eggs were cultured to day 2 and transferred to female mice. Finally, positive mice with a 20 bp frameshift mutation in the second exon of the *TBX15* gene were obtained. Positive mice with the second exon of the *PAX1* gene completely knocked out were also obtained. The discrimination criterion was the presence of a 20 bp frameshift mutation in the second exon of the *TBX15* gene and the second exon of the *PAX1* gene. The mice were raised in a pathogen-free environment and bred according to SPF animal breeding standards. The ambient temperature was 20-25 °C, and humidity 40% -70%. The F0 generation-positive mice were mated with wild-type C57 mice to obtain the heterozygous F1 generation mice.

The heterozygous F1 mice were then mated with a female-to-male ratio of 2:1 to produce the homozygous F2 generation mice. The wild type, heterozygous, and homozygous mice were used for phenotyping when sexually mature at 9 weeks. The use of laboratory animals (SYXK 2019-0022) was licensed by the Beijing Municipal Science and Technology Commission.

### Genotyping

Mice tail tissue of 0.3 cm^3^ was cut into a 1.5 ml centrifuge tube, then added with 98 μL mouse tail lysate and 2 μL proteinase K before incubated in a metal bath at 55 °C for 30 min for full lysis. After that, the proteinase K was inactivated by placing the tube in a 95 °C metal bath for 10 min. The mixture was centrifuged at 12,000 r/min for 5 min, and the supernatant was collected as the DNA template for genotyping. The *Tbx15* and

*Pax1* primers with sequences provided in Table S2 were used for PCR amplification in a 50 μL mixture of 25 μL 2 × Taq Plus Master Mix (Dye Plus), 2 μL upstream primer (concentration: 10 μM), 2 μL downstream primer (concentration: 10 μM), 2 μL DNA template, and 19 μL ultrapure water. The PCR procedures for both gene fragments were as follows: (1) Denaturation at 94 °C for 5 min; (2) Denaturation at 94 °C for 30 s; (3) Annealing at 60 °C for 30 s; with different extension time for (4) *Tbx15* at 72 °C for 2 min, and *Pax1* at 72 °C for 75 s; followed by a final (5) extension at 72 °C for 7 min, steps (2) to (4) were repeated 35 times. Electrophoresis was performed on a 2% agarose gel with a voltage of 125 V and a current strength of 400 A for 30 min. The genotypes were identified with gel electrophoresis and Sanger sequencing. The mouse genotype identification kit was purchased from Nanjing Nuoweizan Biotechnology Co., Ltd. The PCR primers were synthesized by Meiji Biotechnology Co., Ltd.

### Phenotyping

The 9-week-old mice were weighed with an electronic scale and sacrificed by cervical dislocation. The hair of the mice was removed with a razor and Weiting depilatory cream. The body length and limb length of the hairless mice were measured with a ruler. A HandySCAN BLACK scanner was used to scan the hairless mice for the 3D models of the mice. The resolution of the 3D scanner was 0.3 mm, and the scanning accuracy was 0.03 mm. Each point of the model has three-dimensional coordinates, which were used for the subsequent distance calculations. A cup with 135.71 mm in height and 84.55 mm in diameter was scanned to verify the accuracy of the 3D measurement. The 3D model of the cup showed a height of 135.70 mm and a diameter of 84.58 mm, which confirms the accuracy of the 3D measurement (S3 Fig).

Based on the facial features of mice, 17 anatomical feature points were identified to measure the difference in facial morphology (Fig3A-3B). Quality control was performed, and mice with unclear landmarks were rescanned. The images were analyzed by manually extracting landmarks coordinates using the wrap software. Finally, three-dimensional coordinates of 17 landmarks were obtained. After Generalized Procrustes Analysis (GPA), a total of 136 Euclidean distances between all pairs of the 17 landmarks were derived. Euclidean distances as facial phenotypes were used in the subsequent analyses.

### Protein sequence analysis

The Tbx15 and Pax1 protein sequences were obtained from the UniProt database [37,38]. Multiple alignments of amino acid sequences were performed using the DNAMAN software. The analysis of the protein conserved domains was performed using PfamScan software[39].

### Gene expression analysis

RNA-seq data from the study of Prescott et al., GTEx and ENCODE were used to compare the differences in gene expression levels between these two genes in CNCCs and in other types of cells.

### Statistical analyses

(1) 10-week-old heterozygous mice were mated with a female-to-male ratio of 2 : 1, and birth rate of the mice with different genotypes was analyzed using the Student’s t-test analysis. The body weight, body length, and limb length of the mice were analyzed with multiple linear regression analysis.

(2) The association between the genotypes and the phenotypes was analyzed with a general linear regression model (GLM). The genotypes were assigned according to an additive model, which assumes that the number of 20 bp frameshift mutations in the second exon of the *Tbx15* gene and the number of the deletion in the second exon of the *Pax1* gene have a cumulative effect. To correct phenotypic differences resulting from sex, a general linear regression model was used based on the assumptions of the additive model, as shown in Formula (1):

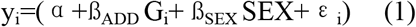

where yi represents a facial characteristic, α represents a fixed effect, ßADD represents the effect of the genotype on facial phenotype, ßSEX represents the gender effect on facial phenotype, εi represents the residual error, Gi represents the genotype, SEX represents gender. The genotype and sex assignment methods are shown in Formulas (2) and (3), respectively.

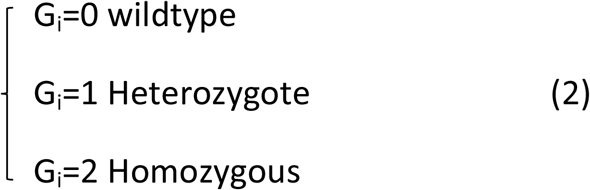

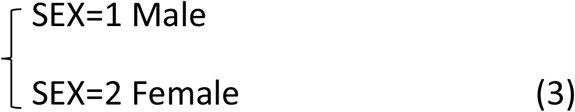

(3) To adjust for the multiple testing of multiple phenotypes, the Bonferroni method was used to correct the effective number of independent variables, which was estimated using the Matrix Spectral Decomposition (matSpD) method [40]

(4) Data analysis and statistical inference were performed using the R 3.5.1 software (https://www.r-project.org/) and the GraphPad Prism 7 software (https://www.graphpad.com/scientific-software/prism/).

## Acknowledgements

This project was supported by the Strategic Priority Research Program of Chinese Academy of Sciences (Grant No. XDB38010400), Shanghai Municipal Science and Technology Major Project (Grant No. 2017SHZDZX01), the Strategic Priority Research Program of Chinese Academy of Sciences (Grant No. XDC01000000), China scholarship council (PhD Fellowship). MK and ZX were supported by Erasmus University Medical Center Rotterdam.

## Author contributions

Conceptualization: Fan Liu, Lei Liu Data curation: Fan Liu, Yu Qian

Formal analysis: Yu Qian, Ziyi Xiong,Yi Li, Haibo Zhou

Funding acquisition: Fan Liu, Manfred Kayser

Investigation: Yu Qian, Ziyi Xiong, Yi Li, Haibo Zhou

Project administration: Fan Liu, Yu Qian, Lei Liu

Resources: Fan Liu, Lei Liu

Supervision: Fan Liu, Yu Qian, Yi Li

Validation: Yu Qian, Yi Li

Writing – original draft: Yu Qian

Writing – review & editing: Fan Liu, Manfred Kayser

## Figure legend

**S1 Fig:Effects of *Pax1* on mice weight, body length, fore and hind limb length.**

**S2 Fig: Amino acid sequence alignments of the *TBX15* and *PAX1* in Homo sapiens and Mus musculus.**

**S3 Fig: 3D measurement example.**

**S1 Table. The sgRNA sequences. S2 Table. The primer sequences.**

**S3 Table. Characteristics of the mice.**

**S4 Table. Characteristics of 136 facial phenotypes in mice.**

**S5 Table. Effects of sex to 136 facial phenotypes in mice.**

**S6 Table. Phenotype correlations between all facial phenotypes in *Tbx15* mice.**

**S7 Table. Effects of *Tbx15* knockout on 136 facial phenotypes in mice.**

**S8 Table. Effects of *Pax1* knockout on 136 facial phenotypes in mice.**

**S9 Table. Difference significance of normalized RNA-seq VST values in each of 2 genes between CNCCs and all 49 types of cells.**

